# Serum-free medium increases the replication rate of the avian coronavirus infectious bronchitis virus in chicken embryo kidney cells

**DOI:** 10.1101/2021.04.30.442100

**Authors:** José A. Quinteros, Glenn F. Browning, Amir H. Noormohammadi, Mark A. Stevenson, Mauricio J. C. Coppo, Carlos A. Loncoman, Nino Ficorilli, Sang-Won Lee, Andrés Diaz-Méndez

**Affiliations:** Asia-Pacific Centre for Animal Health, Melbourne Veterinary School, Faculty of Veterinary and Agricultural Sciences, The University of Melbourne, Parkville, 3010, Victoria, Australia; College of Veterinary Medicine, Konkuk University, 120 Neungdong-ro, Gwangjin-gu, Seoul 143-701, Republic of Korea; Asia-Pacific Centre for Animal Health, Faculty of Veterinary and Agricultural Sciences, The University of Melbourne, Werribee, 3030, Victoria, Australia

**Keywords:** IBV, cell culture, viral replication rate, FBS

## Abstract

Infectious bronchitis virus (IBV), an avian coronavirus, can be isolated and cultured in tracheal organ cultures (TOCs), embryonated eggs and cell cultures. TOCs and embryonated eggs are commonly used for viral isolation but use of these is laborious and expensive. Cell cultures have been used only with IBV strains that have previously been adapted to grow under laboratory conditions, and not for primary isolation. Previous studies using the coronavirus porcine epidemic diarrhoea virus (PEDV) have suggested that foetal bovine serum (FBS), a common component of cell culture media, can inhibit the adsorption of coronaviruses onto the host cell membrane receptors. In the present study, the replication of IBV in primary chicken embryo kidney (CEK) cell cultures and the Leghorn hepatocellular carcinoma (LMH) cell line was examined using two different cell culture media, one containing FBS and the other containing yeast extract (YE). A reverse transcriptase quantitative polymerase chain reaction (RT-qPCR) assay was used to quantify viral RNA copies in cell lysates. The highest concentrations of viral genomes were observed when the cell culture medium did not contain FBS. Examination of the infectivity of virus grown in CEK cell cultures was examined by titration in embryonated chicken eggs, demonstrating that the cell lysate from CEK cell cultures in medium without FBS contained a higher median embryo infectious dose (EID_50_) than that from CEK cell cultures in medium containing FBS. These results suggest that improved replication of IBV in cell cultures can be achieved by the omission of FBS from the cell culture medium. This may enhance the potential for production of vaccines in cell culture and facilitate the isolation of emergent IBV strains in cell cultures.

## Introduction

Infectious bronchitis virus is generally difficult to isolate and propagate in cell cultures. The preferred substrates used are the allantoic cavity of embryonated eggs or tracheal organ cultures [1, 2]. In order to propagate infectious bronchitis virus in cell cultures, serial blind passages are required until the strain is cell-culture adapted and a characteristic cytopathic effect is observed, normally the presence of rounded cells detached from the cell monolayer and/or formation of syncytia [1, 2].

It has been reported that foetal bovine serum (FBS), a component of the growth and maintenance media routinely used in cell cultures, can interfere with the replication of porcine epidemic diarrhoea coronavirus in cell cultures [3] and human coronavirus OC43 [4].

In contrast, the presence of trypsin in cell culture media may enhance the replication of coronaviruses in cell cultures [3], as the S protein must be post-transcriptionally cleaved into its two sub-units, S1 and S2, prior to invasion of the host cell. The cleavage of the S protein is effected by serine proteases, a group of enzymes that includes trypsin [5, 6]. This cleavage allows the S2 subunit to induce fusion of the envelope of the virion with the host cell membrane [7-9]. An additional cleavage site has been described within the S2 subunit (S2’), and cleavage at this site is also necessary for membrane fusion and infection [8].

In the study reported here, a series of experiments were conducted to examine the effect of FBS and trypsin on the replication of IBV in primary chicken embryo kidney (CEK) and Leghorn hepatocellular carcinoma (LMH) cell cultures. LMH cells have been used previously as a substrate for culture of the highly laboratory adapted Beaudette strain of IBV [10]. Replication of the IBV vaccine strain VicS was measured using a reverse transcriptase quantitative polymerase chain reaction (RT-qPCR) assay and the results obtained using this assay were compared to titres determined by inoculating embryonated hen eggs.

## Materials and Methods

### Chicken embryo kidney (CEK) and Leghorn hepatocellular carcinoma (LMH) cells

Kidney cells were prepared from specific-pathogen-free (SPF) chicken embryos that had been incubated until they reached day 17-18 of incubation. Embryonated eggs were transferred to a cold room for at least 4 hours before being extracted from the eggs to reduce sensitivity. Eggs were then placed inside a Class II biological cabinet, the egg shell removed above the air sac, and the embryo removed from the egg, placed in a Petri dish and decapitated, as approved by the University of Melbourne Animal Ethics Committee (Approval Number 1513567). Embryos were dissected and the kidneys removed and placed into a 50 ml tube containing sterile phosphate-buffered saline (PBS). The tube was inverted several times to mix the content, the PBS was removed, and fresh PBS was added. This washing procedure was repeated 3 times. Kidney tissue was then minced with a scalpel blade into fine pieces and transferred into 100 ml of 0.125% trypsin-Versene. The suspension was mixed and stored overnight at 4°C. The following day, the suspension was shaken for 3 minutes and the tissue then filtered through a piece of sterile gauze over a sterile beaker to separate the remaining connective and adipose tissue from the kidney cells. The cell suspension was transferred into 50 ml tubes and centrifuged at 524 × g for 5 minutes at 4°C. The supernatant was removed, and the cells were resuspended in 10 ml of cold PBS. A 30 μl volume of the suspension was mixed with 30 μl of trypan blue and a viable cell count was performed using a haemocytometer. Sufficient growth medium (Dulbecco’s modified Eagle’s medium [DMEM] supplemented with 10% v/v FBS, gentamicin [0.05 mg/ml], amoxicillin [0.05 mg/ml] and amphotericin B [0.005 mg/ml]) was added to the suspension to achieve a viable cell concentration of approximately 1 × 10^6^ cells per ml.

The LMH cell monolayers were cultured in a growth medium (GM) of DMEM supplemented with amphotericin B (0.005 mg/ml), gentamicin (0.05 mg/ml), co-trimoxazole (0.01 mg sulfamethoxazole/ml and 0.002 mg trimethoprim/ml), 10% v/v FBS and 10 mM HEPES (4-(2-hydroxyethyl)-1-piperazineethanesulfonic acid, pH 7.7).

CEK and LMH cells were seeded into 6-well plates that had previously been coated with a sterile solution of 0.1% gelatine in PBS (pH of 7.6) and incubated overnight at 4°C. Before cell seeding, the gelatine was removed from the wells, and 3 × 10^6^ cells (3 ml of cell suspension) were inoculated into each well. The plates were then incubated at 37°C in 5% CO_2_. After 24 hours, the cell monolayers had reached approximately 70% confluence and were inoculated with virus.

### Preparation of inocula

In the CEK cell culture experiment, two different inocula were used. One inoculum was prepared using allantoic fluid (AF) from embryonated eggs infected with the IBV strain VicS-v. The AF contained 2 × 10^5^ median embryo infectious doses (EID_50_) of VicS-v/ml and was diluted one hundred-fold to yield a final titre of 2 × 10^3^ EID_50_/ml. One half of this diluted AF was used as untreated inoculum, while the other half was treated with trypsin before inoculation. Both inocula were prepared in DMEM supplemented with gentamicin (0.05 mg/ml), amoxicillin (0.05 mg/ml) and amphotericin B (0.005 mg/ml). This same medium, without the addition of virus, was used in the negative control groups (mock-inoculum). The inoculum supplemented with 15 μl of 1% trypsin was referred as the trypsinised inoculum (TI), while the unsupplemented inoculum was referred as the non-trypsinised inoculum (NTI). Neither inoculum contained FBS.

In the LMH cell culture experiment, two different inocula were used. The first inoculum was the same allantoic fluid containing VicS-v as was used in the CEK experiment (without the addition of trypsin). The other inoculum was the cell lysate from CEK cell cultures previously inoculated with VicS-v. Details about this cell culture lysate are provided below.

### Preparation of maintenance media

In the CEK cell culture experiment, four different maintenance media were prepared. All contained DMEM with gentamicin (0.05 mg/ml), amoxicillin (0.05 mg/ml) and amphotericin B (0.005 mg/ml). The medium supplemented with 10 μg of trypsin/ml was referred as the trypsinised medium (TM) and the medium without trypsin was referred as non-trypsinised medium (NTM). Half of each of these media was supplemented with 1% v/v FBS (CSL) and the other half of each was supplemented with 0.02% YE (Oxoid). Thus, the media prepared were FBS with trypsin (FBS TM), FBS without trypsin (FBS NTM), YE with trypsin (YE TM), and YE without trypsin (YE NTM). The experiment contained three biological replicates of each treatment and is summarised in Fig 1. The 1% YE used to prepare the media was diluted just before the preparation of the media and filter-purified though a 0.45 μm syringe filter.

**Fig 1.**
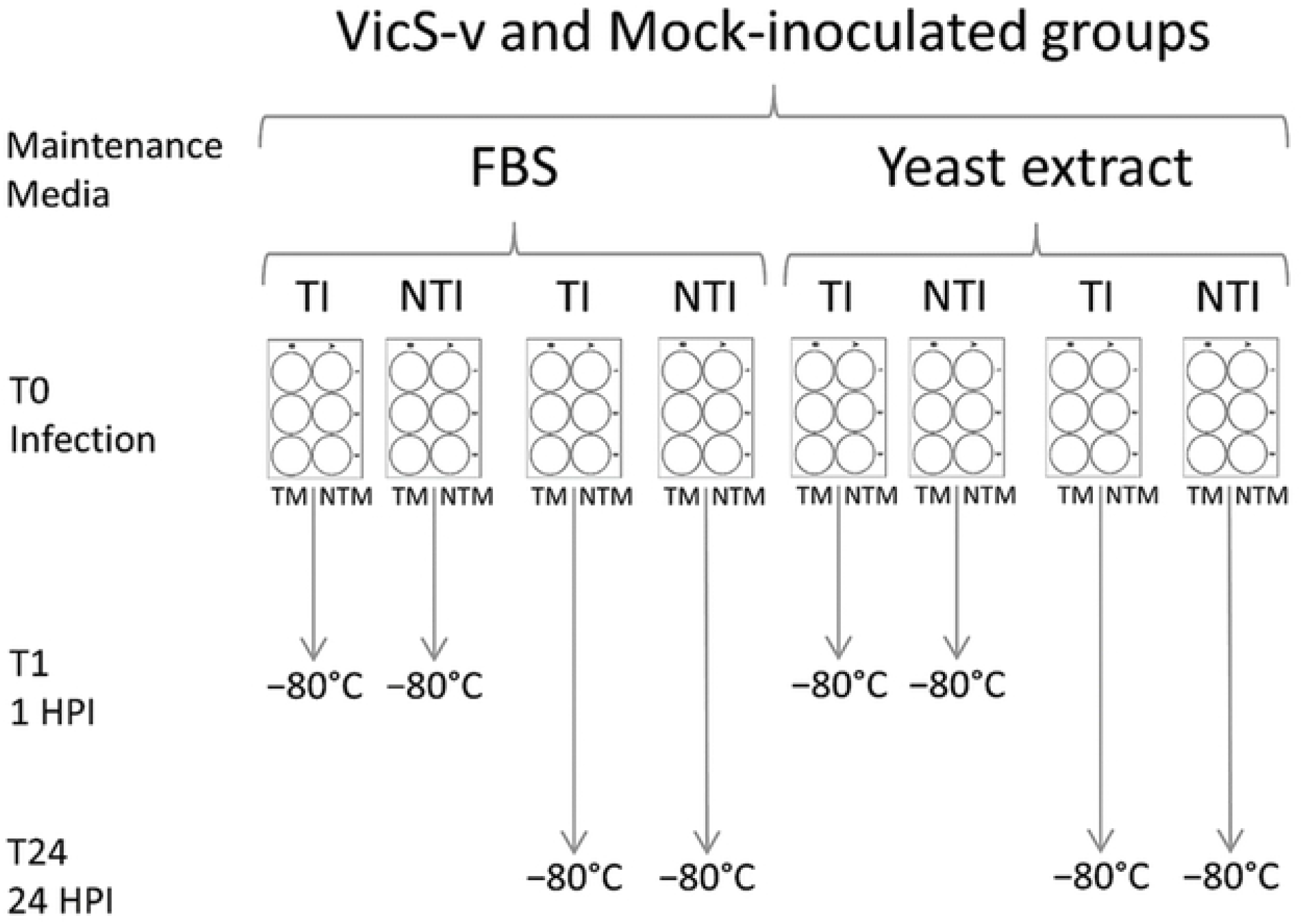
Experimental design of the CEK cell culture study, using inocula with and without trypsin (TI and NTI) and maintenance media with and without trypsin (TM and NTM), and with FBS or YE. T1 was at 1 hour post infection (HPI), 1 h after the cell monolayer was inoculated; T24 was 24 HPI, 24 h after the cell monolayer was inoculated.

In the LMH cell culture experiment, the maintenance medium was DMEM containing gentamicin (0.05 mg/ml), amoxicillin (0.05 mg/ml) and amphotericin B (0.005 mg/ml). Half of the medium was supplemented with 1% v/v FBS and the other half with 0.02% YE.

### Cell culture infection and incubation

Cells were inoculated, starting with the mock inoculum, followed by the AF or the cell lysate containing VicS-v. The growth medium was removed, and each well was washed with 2-4 ml of PBS. A 400 μl volume of inoculum was then added to each well. Plates were then transferred into a 5% CO_2_ incubator at 37°C and manually rocked every 15 min for 1 hour. The inocula were then removed and the wells washed with 2-4 ml of PBS. Finally, 3 ml of maintenance medium was added into each well (Fig 1). One 6-well plate was immediately transferred to a freezer at −80°C. The rest of the plates were incubated for 24 hours, and then transferred to a freezer at −80°C. For the LMH cell experiment, an additional set of plates were incubated for 72 hours, then stored at −80°C.

### Cell lysis and collection of the lysates

In order to lyse the cells, the 6-well plates were transferred from −80°C to a warm room at 37°C until the content was completely thawed, and then transferred back to the freezer at -80°C. This was repeated three times. The total content of each well was transferred into individual 10 ml tubes and stored at −80°C.

### RNA extraction

The RNA was extracted using the PureLink Pro 96 viral RNA/DNA purification kit (Invitrogen, USA). The extraction was performed in a QIAGEN-Corbett X-tractor Gene automated DNA/RNA extractor (QIAGEN, Australia) using the default settings for RNA extraction. After thawing and mixing the cell lysates, a 100 μl sample of each lysate was pipetted into a 96-well plate and nucleic acid extraction was performed. The final elution step was performed using 70 μl of elution buffer. The VicS-v- and mock-inoculated samples were processed in separate 96-well plates.

### Reverse transcriptase quantitative polymerase chain reaction (RT-qPCR) assays

The RT-qPCR assays were performed in two steps. The reverse transcription step was performed using Superscript III reverse transcriptase (Invitrogen), following the manufacturer’s instructions. A 5 μl sample of RNA was first added to the master mix, each aliquot of which contained 1 μl of random hexamers (approximately 1.8 ng), 1 μl of dNTPs (0.5 mM) and nuclease-free water to a total volume of 13 μl. The mixture was incubated in a thermocycler at 65°C for 5 min, and the reactions were then immediately transferred onto ice. A second master mix aliquot containing 4 μl of 5× first-strand buffer (50 mM Tris-HCl (pH 8.3), 75 mM KCl, 3 mM MgCl2; Invitrogen), 1 μl of dithiothreitol (DTT, 5 mM; Invitrogen), 1 μl of RNase-OUT (40 U; Invitrogen) and 1 μl of Superscript III reverse transcriptase (200 U; Invitrogen) was added to each reaction, and the reactions transferred to a thermocycler and incubated at 55°C for 1 h, then at 25°C for 5 min and finally at 70°C for 15 min. The reactions were then frozen at −20°C.

The cDNA was then assayed using two different qPCRs. The first qPCR targeted the gene of interest (GOI), the N protein-3’UTR region of the IBV genome, and the second qPCR targeted a housekeeping gene (HKG), the mRNA of the chicken GAPDH gene (Table 1). For the GOI qPCR, a master mix was prepared consisting of 5 μl of 5× colourless buffer, 2 μl of each primer (2 μM each), 2 μl of MgCl2 (1 mM), 4 μl of dNTPs (1.25 mM of each), 2 μl of SYTO 9 green fluorescent nucleic acid stain (8 μM), 0.25 μl GoTaq polymerase (1.25 U; Promega) and 5 μl of cDNA. For the HKG qPCR, a master mix was prepared consisting of 5 μl of 5× colourless buffer, 1.25 μl of each primer (0.5 μM), 2 μl of MgCl2 (1 mM), 4 μl of dNTPs (1.25 mM of each), 2 μl of SYTO 9 green fluorescent nucleic acid stain (8 μM), 0.25 μl GoTaq polymerase (1.25 U, Promega) and 5 μl of cDNA. The HKG primers used in this experiment targeted the GAPDH mRNA exclusively, not the DNA, as they each bridge exon-exon junctions.

**Table 1.**
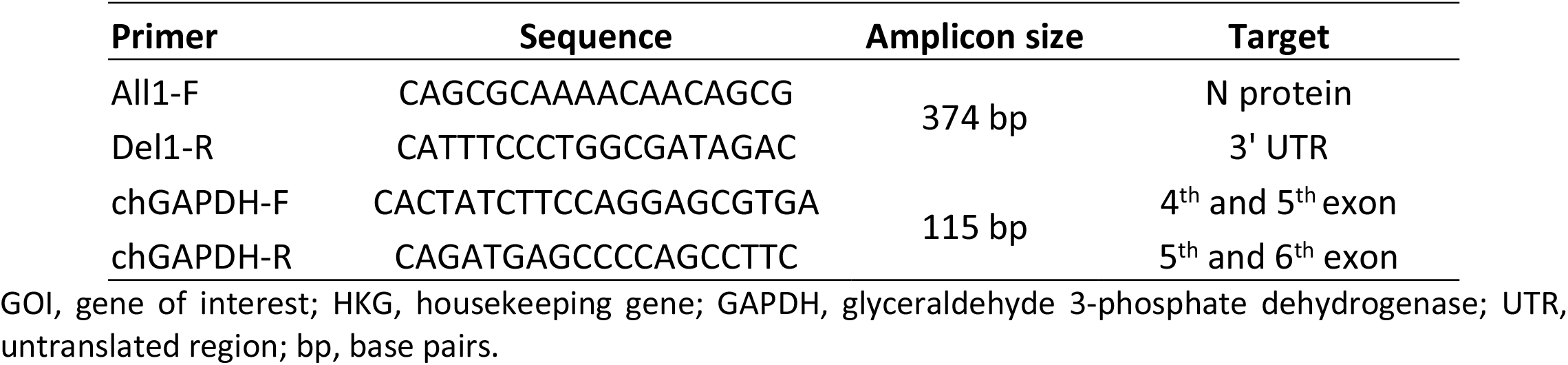
Primers used for the amplification of the GOI (IBV) and the HKG (chicken GAPDH).

The 2^−ΔΔCt^ values for each sample were calculated from the Ct values for the GOI and HKG qPCRs, and the Ct values for the same genes for the corresponding qPCR positive controls. The 2^−ΔΔCt^ values were log_2_ transformed and the values at 1 hour post infection (HPI), 24 HPI and 72 HPI (LMH experiment) were compared for each medium and treatment using Tukey’s honestly significant difference (HSD) test. The log_2_ transformation allowed comparison of two-fold increases/decreases in the number of copies of RNA, and has been used previously for comparisons of gene expression, for microarray analysis, and for viral load comparisons [11-14]. Similar methods using the log_2_ transformation of ΔCt or ΔΔCt values have been described previously for quantitation and comparison of gene expression in terms of mRNA quantities and for comparisons of viral loads [14-17].

### Titration of the CEK supernatants

Viral titrations were performed on the cell lysates obtained at 1 and 24 HPI from the CEK cell cultures with and without FBS, and with the NTINTM medium. The triplicates for each treatment were thawed, and 200 μl samples of each cell lysate triplicate were mixed and homogenised in order to obtain a single sample for each treatment at each time point. Four cell lysates were tested: FBS NTINTM at 1 HPI, FBS NTINTM at 24 HPI, YE NTINTM at 1 HPI and YE NTINTM at 24 HPI. The titration was performed following the protocol of the United States Department of Agriculture (USDA) Center for Veterinary Biologics Supplemental Assay Method 411 “Titration of Newcastle Disease Vaccine, Infectious Bronchitis Vaccine and combination of Newcastle Disease/Infectious Bronchitis Vaccine in Chicken Embryos” [18]. The samples were aseptically inoculated into the allantoic cavity of 9-day-old embryonated SPF eggs. The inocula used were an undiluted sample and samples from four tenfold serial dilutions (that is, up to 10^−4^), with 5 embryonated eggs inoculated per dilution, and 4 positive control eggs inoculated with serial dilutions of allantoic fluid containing the IBV strain VicS-v, and 4 negative control eggs mock inoculated with the dilution medium used to prepare the inocula (low-bicarbonate DMEM supplemented with gentamicin [0.05 mg/ml], amphotericin B [0.005 mg/ml] and 1% v/v FBS). The eggs were inspected daily for 7 days and any embryos that died over the first 24 hours post inoculation were discarded as non-specific deaths. At 7 days post inoculation, all the embryos were euthanised and individually inspected. All those embryos exhibiting pathological changes typical of infection with IBV (dwarfing, curling, clubbed down) were considered positive. During analysis, high-resolution photographs were taken and then sent to two additional assessors (blind evaluators). The titre was calculated using the data from the three assessors using the Spearman-Kärber method [5]. These three independent titrations were used for the statistical comparisons of the titres.

## Results

### Viral replication rate was higher in media without FBS

In the experiment using CEK cell cultures, the log_2_ transformed 2^−ΔΔCt^ values at 1 HPI and 24 HPI did not differ significantly for any of the treatments containing FBS (Table 2). However, there was a significant difference between 1 HPI and 24 HPI for all the treatments in which the FBS in the medium was replaced with YE (Table 2). The larger increments in the log_2_ 2^−ΔΔCt^ values were seen in the YE NTINTM (5.67) and YE TINTM (5.11) treatments (Table 2).

**Table 2.**
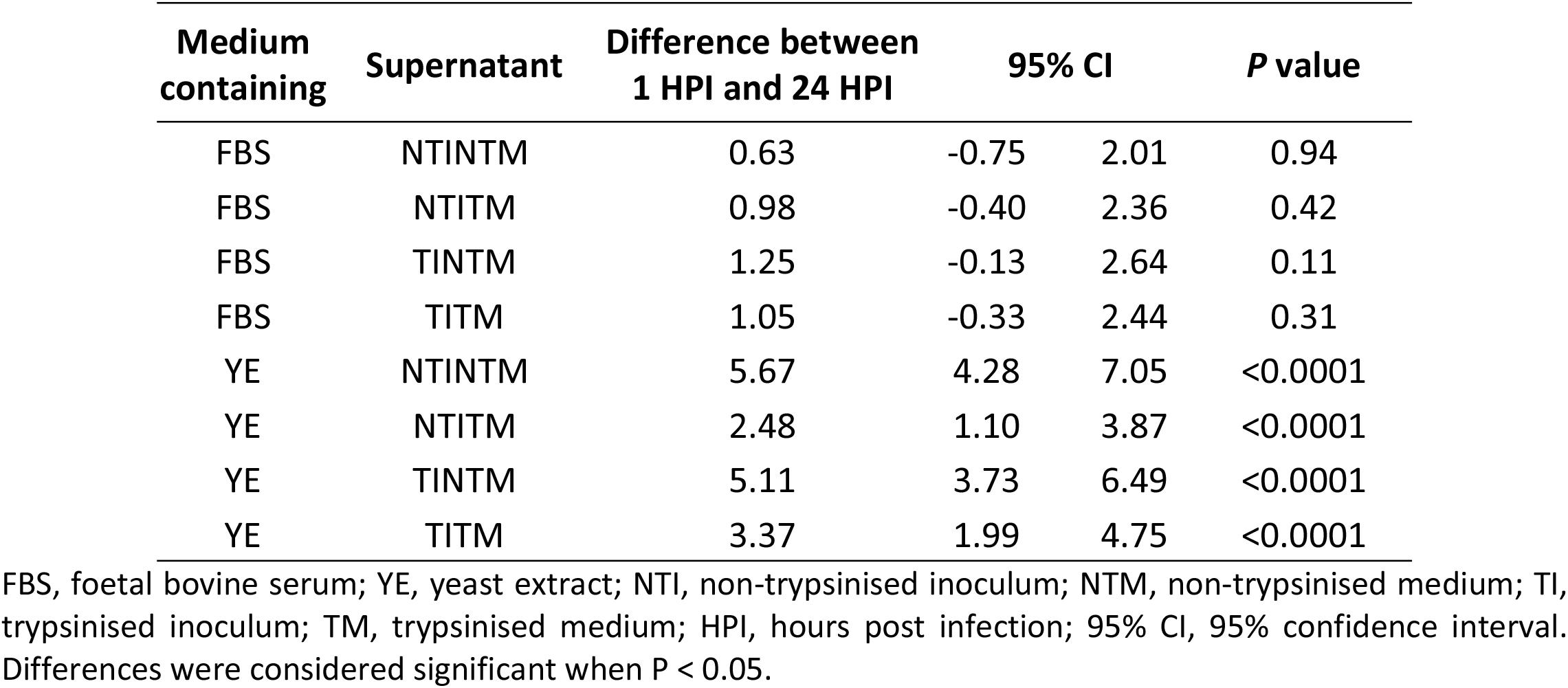
Differences in IBV VicS-v viral RNA concentrations between 1 and 24 hours post infection of CEK cell cultures expressed as the difference between the log_2_ transformed 2^-ΔΔCt^ values.

In the experiment using LMH cells, the effect was similar. There were no differences between 1, 24 and 72 HPI in any of the treatments containing FBS. The biggest increase in viral concentration was seen in those cultures that were infected using the cell lysate from a CEK cell culture with the treatment NTINTM (Table 3). There was no increase in viral concentration between 24 and 72 HPI for any of the cultures.

**Table 3.** Differences in IBV VicS-v viral RNA concentrations between 1 and 24 hours post infection, and between 24 and 72 hours post infection of LMH cell cultures, expressed as the differences between the log_2_ transformed -2^ΔΔCt^ values.

| Media Containing | Inoculum | Difference between 1 HPI and 24 HPI | P value | Difference between 24 HPI and 72 HPI | P value |
| --- | --- | --- | --- | --- | --- |
| FBS | AF | 2.09 | >0.9999 | 2.70 | >0.9999 |
| FBS | CCL | -0.77 | >0.9999 | 1.79 | >0.9999 |
| YE | AF | 4.24 | 0.42 | 0.04 | >0.9999 |
| YE | CCL | 5.54 | 0.12 | -0.29 | >0.9999 |

FBS, foetal bovine serum; YE, yeast extract; AF, allantoic fluid; CCL, cell culture lysate; HPI, hours post infection. Differences were considered significant when P < 0.05.

### Titres of IBV VicS-v cultured in CEK cells

The greatest differences in viral RNA concentrations between 1 and 24 HPI were detected when the NT inoculum and NT medium (NTINTM) were used, and when the medium was supplemented with YE rather than FBS (Table 2). Therefore, lysates from NTINTM cultures were used to compare titres of viable virus obtained using media containing FBS or YE. As shown in Table 4, the titres at 1 HPI were similar in the cultures with FBS (1.08 × 10^1^ EID_50_ per ml) or without FBS (1.47 × 10^1^ EID_50_ per ml). However, at 24 HPI the viral titres in the cultures with FBS had increased to 0.68 × 10^3^ EID_50_ per ml, while in the cultures with YE (without FBS) the titres had increased to 5.01 × 10^3^ EID_50_ per ml (Table 4). The geometric mean titres were calculated using the assessments of the three independent observers. The titres were compared using a one-way ANOVA and Tukey’s multiple comparison test. As shown in Table 4, the titres at 1 HPI did not differ significantly between cultures with or without FBS (*P* > 0.05). The difference in titre between 1 and 24 HPI was significant in cultures in maintenance medium with FBS (*P* < 0.05) or with YE (*P* < 0.0001), and the titres at 24 HPI differed significantly between cultures in maintenance medium with FBS and those in maintenance medium with YE (P < 0.0001), with higher titres obtained in cultures that did not contain FBS.

**Table 4.** Viral titres in the lysates of CEK cell cultures in maintenance medium with FBS or YE at different time points (1 HPI and 24 HPI).

| Cell lysate | Titre (EID<sub>50</sub>/ml) |
| --- | --- |
| FBS 1 HPI | $1.08 \times 10^1$ <sup>a</sup> |
| FBS 24 HPI | $0.68 \times 10^3$ <sup>b</sup> |
| YE 1 HPI | $1.47 \times 10^1$ <sup>a</sup> |
| YE 24 HPI | $5.01 \times 10^3$ <sup>c</sup> |

Titres marked with the same superscript letter did not differ significantly ( $P > 0.05$ )

## Discussion

This study was conducted to determine whether omitting FBS from the maintenance medium and/or including trypsin in the medium could improve the yields of IBV in cell cultures. Different treatment combinations were assessed, with IBV detected by RT-qPCR. The rationale behind including the estimation of the GAPDH mRNA copies during the RT-qPCR was to normalise the amount of viral RNA to the number of cells collected from the cell cultures and lysed for measurement of viral titres. Titrations in embryonated eggs were also used to determine the concentration of viable virus in the same cultures. We found that a 24-hour culture period was sufficient to detect an increase in IBV replication. A previous study using SARS-CoV cultured in cell monolayers reported that at 10 hours post infection it was possible to identify a significant number of fully formed virions [19].

There was a significant increase in the concentrations of viral genome in the CEK cell cultures containing FBS at 24 HPI (Table 4). However, viral replication was greater in the cultures from which FBS was omitted from the cell culture maintenance medium. In previous work [20], the inclusion of FBS at 7% in the agar overlay used for plaque counting was found to reduce the titres of several strains of IBV (97, SE 17, JMK, 13721, 46 and 41) by 10- to 100-fold. This reduction in the titre was attributed to non-specific inhibition.

The greatest increase in viral RNA concentrations at 24 HPI was seen in cell cultures in which neither FBS nor trypsin were added to the maintenance medium (NTINTM). It is possible that the addition of trypsin to the maintenance medium could have an adverse effect on the virus or the cell monolayer.

It seems that the addition of trypsin to the maintenance medium or the inoculum had little effect on the viral replication rate in cell cultures. The differences between 1 HPI and 24 HPI were similar in the NTINTM treatment and the TINTM treatment.

Similar results were seen in LMH cell cultures, although the differences between the time points were not significant. The replication of the virus appeared to be improved when FBS was not included in the maintenance medium. The greatest increase in viral RNA copy concentrations at 24 HPI was seen in the LMH cell cultures infected with IBV strain VicS-v that had previously been cultured in CEK cells, suggesting that the replication of the virus was enhanced by prior CEK passage.

To confirm that the differences in the viral RNA copy concentrations correlated with the titres of viable virus, the cell culture lysates were titrated in embryonated eggs. The greatest increase in viral titre was seen in NTINTM cultures in maintenance medium containing YE rather than FBS. The differences in viral RNA concentrations calculated using RT-qPCR assays only indicate differences in the presence of IBV genome between the different treatments. However, the differences in the titres demonstrated that there were indeed more infective viral particles in the cultures without FBS in the medium than in those with FBS in the medium (Table 4).

While these studies have shown that FBS can interfere with the replication of IBV in CEK cells, and possibly also in LMH cells, the underlying reason for this interference is yet to be elucidated. It has been observed that in vesicular stomatitis virus (VSV), a negative stranded RNA virus, bovine serum albumin, the main component of FBS, can cause depletion of cholesterol in the membrane of the virion, resulting in a decrease in its infectivity [21]. Moreover, It has been shown that bovine serum contains β inhibitors, which interfere with the infectivity and haemagglutination activity of influenza A viruses [22, 23].

It is possible that interference by FBS occurred not during the initial attachment of the virus to the host cells, but rather during the multiple subsequent rounds of replication, when virus was transmitted from cell to cell. This is based on the fact that the media with and without FBS were added one hour after infection of the cell monolayers, so it is possible that the initial virion-receptor interaction had already occurred. Similar interference with viral replication has been reported previously in other viruses, including a decrease in release of Epstein-Barr virus when adult bovine serum was included in the cell culture medium [24]. Further studies examining growth rates of IBV in cell culture media with and without FBS, and examining cell-to-cell spread of virus in these media may help to further elucidate the basis of this interference.

## Acknowledgments

The authors wish to acknowledge the advice of Dr Jagoda Ignjatovic and Ms Denise O’Rourke. This study was supported by the Australian Research Council, the University of Melbourne Graduate Research Scholarship Program and the Government of Chile through the Becas Chile Scholarship Program.

